# Efficient differentiation of human retinal pigment epithelium cells from chemically induced pluripotent stem cells

**DOI:** 10.1101/2024.03.22.586212

**Authors:** Ke Zhang, Yanqiu Wang, Qi An, Hengjing Ji, Defu Wu, Xuri Li, Xuran Dong, Chun Zhang

## Abstract

Human induced pluripotent stem cells (hiPSCs) hold considerable promise for autologous cellular therapies, particularly in ocular disease treatment, because of the low numbers of cells required for transplantation and the non-invasive nature of graft monitoring. However, the use of hiPSCs in ocular disease treatment faces challenges because the production of clinical-grade autologous hiPSCs via genetic techniques remains expensive, time-consuming, and subject to safety concerns. Here, we utilize a recently reported chemical method to derive human chemically induced pluripotent stem cells (hCiPSCs), demonstrating that hiPSC lines can be generated using small molecules in a simple and robust manner. Moreover, we show that these cell lines can be efficiently differentiated into retinal pigment epithelium (RPE) cells, which may aid in the treatment of age-related macular degeneration (AMD). Our study provides a foundation for cost-effective, fast, and safe methods that enable efficient production of autologous retinal cell types for ocular disease treatment.

## Introduction

Pluripotent stem cells, including embryonic stem cells (ESCs) and induced pluripotent stem cells (iPSCs), exhibit unlimited proliferation capacity and can differentiate into diverse functional cell types that are valuable in cell therapy applications^**1**^. Compared with human ESCs (hESCs) derived from human embryos, the production of human iPSCs (hiPSCs) from somatic cells avoids potential ethical concerns and permits generation of autologous iPSCs, enabling personalized cellular therapies without the need for immunosuppressive treatment^**2,3**^. Consequently, the transplantation of autologous iPSC-derived functional cells has emerged as a novel therapeutic strategy for various diseases.

The eye stands out as a major target for hiPSC-based cellular therapies among various targeted organs. Its appeal lies in the lower cell quantity required for therapeutic effects compared to other organs. Additionally, the non-invasive monitoring and direct observation of the engraftment in the eye enable early detection of potential abnormalities, making it a favorable option for therapeutic interventions. Age-related macular degeneration (AMD), the leading cause of central vision loss worldwide, is a key domain of ocular disease research^**4–8**^. The pathophysiology of AMD involves dysfunction and permanent loss of retinal pigment epithelium (RPE) cells, which are essential for maintenance of the subretinal microenvironment. Therefore, the replacement of RPE cells or tissue is regarded as a promising therapeutic strategy for AMD. Notably, over the past decade, RPE cells have emerged as a focus in clinical studies, underscoring their significance in AMD research^**9**^. A clinical study carried out by RIKEN in 2013 attracted attention worldwide; it represented the first clinical trial of autologous hiPSC-RPE cell transplantation, with an emphasis on applying iPSC-based personalized medicine in a clinical setting^**7**^. However, progress in autologous hiPSC-based clinical trials for AMD treatment has faced certain challenges. In RIKEN’s study, Patient 2 did not undergo transplantation because of genetic anomalies present in their hiPSCs^**10**^. Additional obstacles include the time-consuming and expensive nature of genetic methods for production of clinical-grade autologous iPSCs^**11**^. These challenges hinder the large-scale production of autologous functional cells for clinical treatment of retinal diseases, emphasizing the need for technological advancements that can address these practical issues.

Recent advances in the development of chemical methods for iPSC generation have revealed a promising and viable alternative approach^**12–14**^. Theoretically, the use of human chemically induced pluripotent stem cells (hCiPSCs) could avoid safety concerns associated with conventional transgenic techniques. The small molecules utilized in chemical reprogramming do not integrate into the genome^**13,15**^. Furthermore, small molecules are easily manufactured, standardized, and more cost-effective, making them beneficial for large-scale clinical applications^**12–14**^. These benefits indicate that hCiPSC technology can facilitate the production of autologous iPSCs for therapeutic purposes, particularly in clinical settings. To our knowledge, no studies have demonstrated the use of hCiPSCs to generate RPE cells, the most extensively studied therapeutic cell type in clinical research over the past decade.

In this study, we aimed to generate iPSCs through chemical induction and efficiently differentiate them into RPE cells. The hCiPSC-derived RPE cells exhibited distinct RPE characteristics in terms of morphology, gene expression patterns, and functionality, indicating their potential value in future clinical applications.

## Results

### Reprogramming of hADSCs into hCiPSCs

For generation of hCiPSCs from hADSCs, we followed a detailed protocol^**14**^ and utilized the Human iPS Cell Chemical Reprogramming Kit (BeiCell™) to reprogram hADSCs through a three-step induction process. The cells exhibited a series of morphological changes during chemical reprogramming (Figure 1A). After 4 days in the first induction stage, the cells transitioned from a spindle-shaped, fibrous structural morphology to an epithelial cell monolayer. At the end of the second induction stage, a large number of multilayered colonies emerged. During the third induction stage, compact dome-shaped clones were evident.

**Figure 1.**
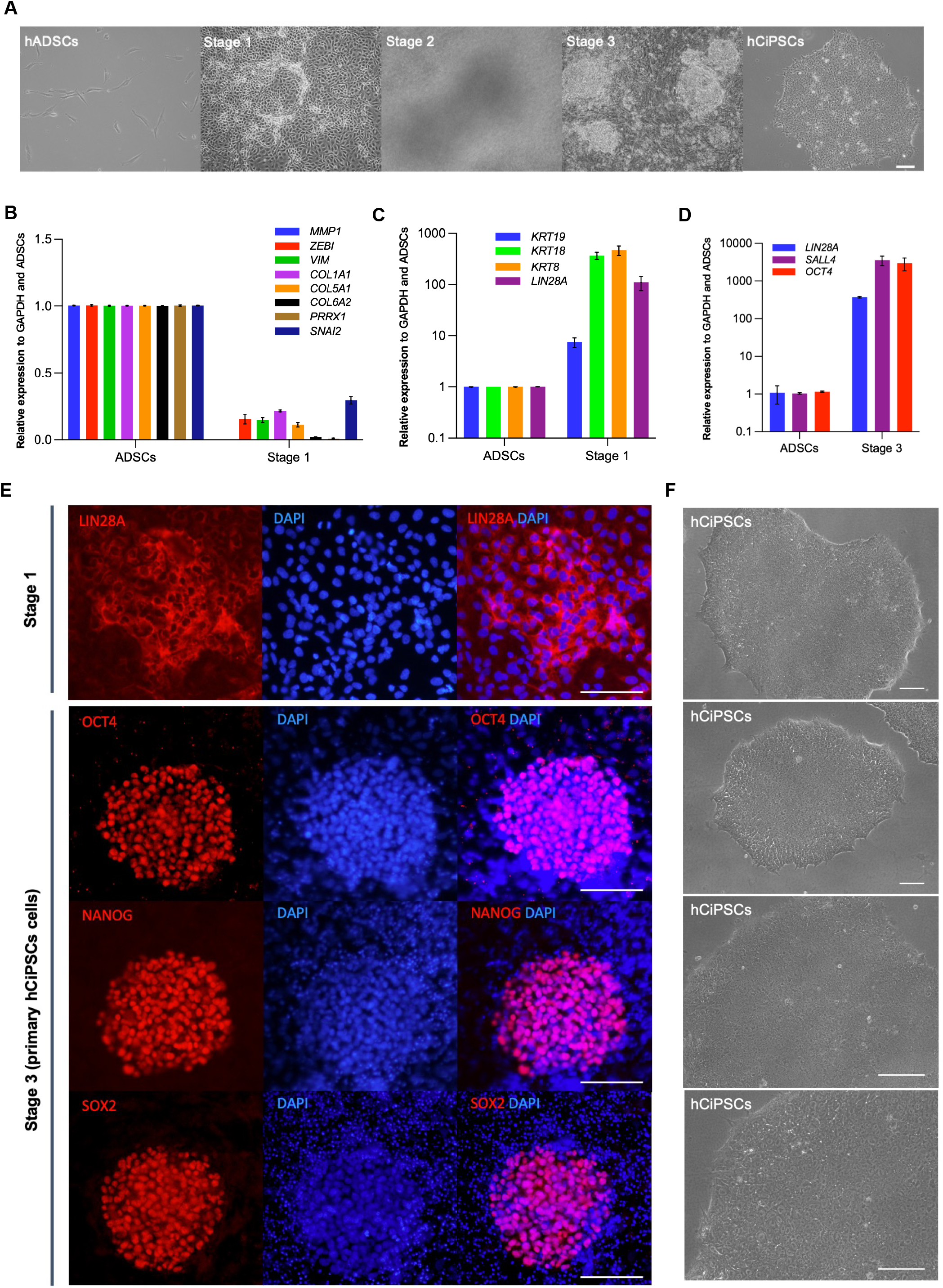
Reprogramming of hADSCs into hCiPSCs. (A) Representative images of initial hADSCs and cells at the end of each stage during hCiPSCs induction. Scale bar, 100μm. (B) Relative expression levels of somatic cell marker genes at the end of the first induction stage, as determined by RT-qPCR. Data are presented as means; n = 3. (C) Relative expression levels of epithelial cell marker genes and *LIN28A* at the end of the first induction stage, as determined by RT-qPCR. Data are presented as means of three independent experiments (n = 3). (D) Relative expression levels of *LIN28A, SALL4*, and *OCT4* at the end of the third induction stage, as determined by RT-qPCR. (E) Immunofluorescence analysis of pluripotency markers at the end of the first (LIN28A) and third (OCT4, NANOG, SOX2) induction stages. Scale bar, 100μm. (F) Representative images of established hCiPSC lines with typical hESC morphologies. Scale bar, 100 μm.

Next, we analyzed fibroblastic and epithelial marker gene expression patterns during the induction process. Reverse transcription (RT)–quantitative polymerase chain reaction (qPCR) analysis showed that the expression levels of fibroblast marker genes (e.g., *MMP1, ZEB1, VIM, COL1A1, COL5A1, COL6A2, PRRX1*, and *SNAI2*) were decreased in treated cells at the end of the first induction stage (Figure 1B). In contrast, epithelial marker genes, such as *KRT19, KRT18*, and *KRT8*, demonstrated substantial upregulation. We also analyzed the expression of *LIN28A*, a key marker of pluripotency that is activated during chemical reprogramming^**16**^ (Figure 1C). In addition to *LIN28A*, two other key pluripotency markers *OCT4* and *SALL4* were expressed at the end of the third induction stage (Figure 1D). We confirmed the expression of these pluripotency factors by immunofluorescence analysis (Figure 1E). Our results indicated that, by the third induction stage, the reprogrammed cells had acquired pluripotency features and became hCiPSCs. Next, we picked 42 colonies and generated 40 hCiPSC lines that could be passaged. Among these 40 cell lines, 12 were randomly selected for further characterization and differentiation experiments. All established cell lines displayed morphologies typical of hESCs (Figure 1F).

### Characterization of hADSC-derived hCiPSCs

During expansion, the established hADSC-derived hCiPSCs exhibited a doubling time similar to the doubling time of hESCs (Figure 2A). After extensive passaging, these cells strongly expressed representative pluripotent cell surface markers (e.g., TRA-1-81 and TRA-1-60, LIN28A), as well as pluripotency transcription factors including OCT4, SOX2, and NANOG (Figures 2B, S1A and S1B). Consistent with these results, RT-qPCR analysis showed that hADSC-derived hCiPSCs expressed multiple pluripotency marker genes at levels comparable to the levels in hESCs (Figures 2C and S1C). Bulk RNA-sequencing analysis of 6 hCiPSC lines revealed that hCiPSCs and hESCs display similar transcriptomic profiles (Figures 2D-2E and S1D), demonstrating that hCiPSCs had successfully exited the somatic state and transitioned to a pluripotent state similar to hESCs (Figure 2F). These results indicated that hADSC-derived hCiPSCs exhibit typical pluripotency features similar to the characteristics of hESCs.

**Figure 2.**
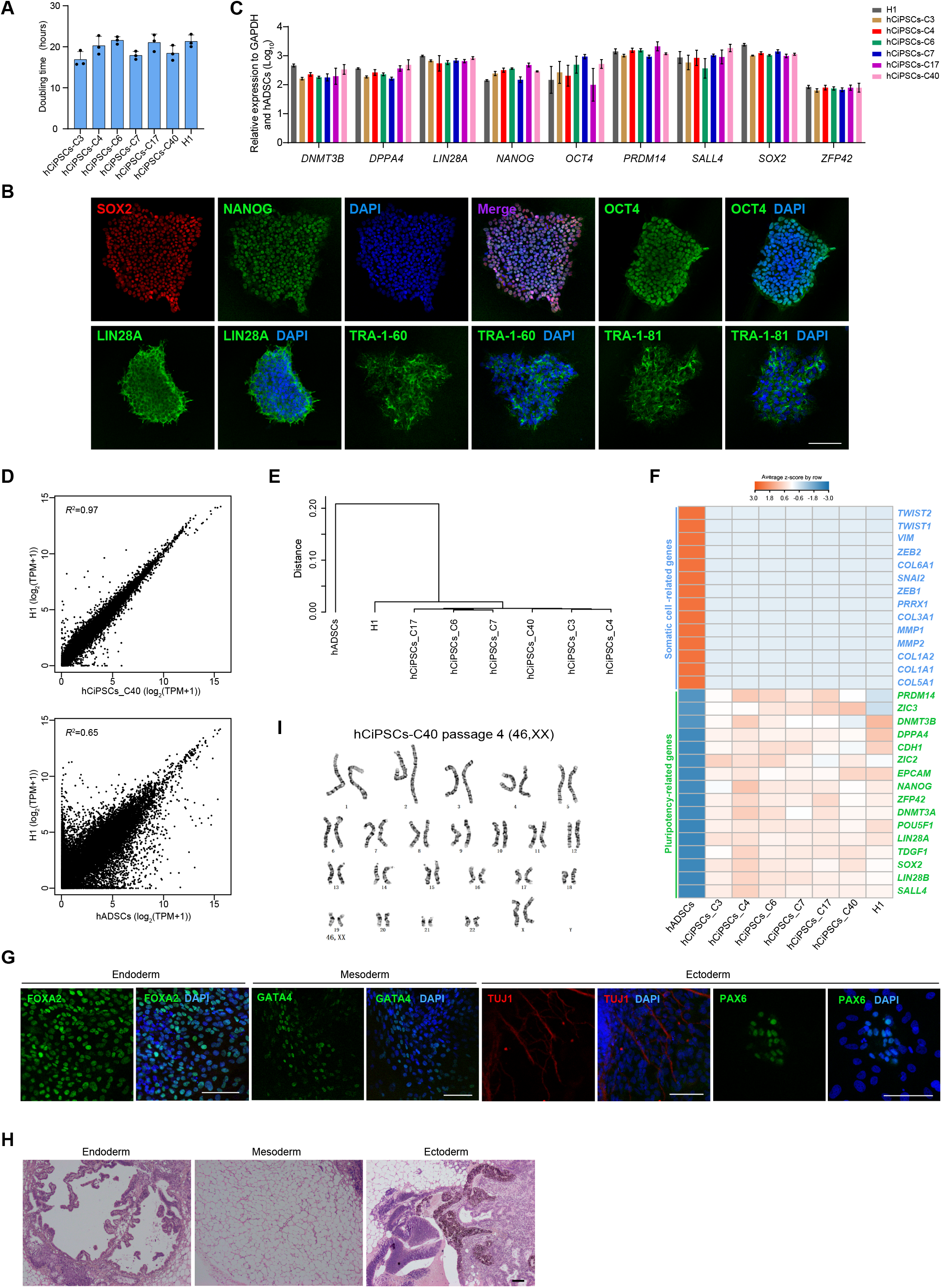
Characterization of hADSC-derived hCiPSCs. (A) Calculated doubling times for hCiPSCs and hESCs. Data are presented as means ± standard deviations (SDs); n = 3. (B) Immunofluorescence staining for pluripotency markers in hADSC-derived hCiPSCs. Scale bar, 100 μm. (C) Relative expression levels of pluripotency marker genes in hESCs (H1) and hADSC-derived hCiPSCs, as determined by RT-qPCR. Data are presented as means ± SDs; n = 3. (D) Scatter plots comparing global transcriptional profiles of hCiPSCs, hESCs (H1), and hADSCs. (E) Hierarchical clustering of the global transcriptional profiles of hCiPSCs, hESCs (H1), and hADSCs. Distances were calculated as 1–Spearman correlation coefficient. (F) Heatmap showing the expression patterns of pluripotency marker genes and somatic cell marker genes in hCiPSCs, hESCs (H1), and hADSCs. (G) Immunofluorescence staining for the three germ layer markers in embryoid bodies derived from hCiPSCs (hCiPSCs-C40, C4). Scale bar, 100 μm. (H) Hematoxylin and eosin staining of hCiPSC-derived teratoma sections (hCiPSCs-C4). Images showing the endoderm, mesoderm, and ectoderm differentiated from a single teratoma. Scale bars, 100 μm. (I) Karyotype analysis showing hADSC-derived hCiPSCs (hCiPSCs-C40) with normal diploid chromosomal content.

Next, we investigated the differentiation potentials of hADSC-derived hCiPSCs through *in vitro* embryoid body differentiation, which showed that spontaneously differentiated cells expressed multiple markers of the three germ layers, such as the endoderm marker FOXA2, mesoderm marker GATA4, and ectoderm markers PAX6 and TUJ1 (Figure 2G). We also conducted *in vivo* teratoma formation assays using hADSC-derived hCiPSCs. Teratomas were formed approximately 1 month after injection into immunodeficient mice. Moreover, hematoxylin and eosin staining analysis showed that hADSC-derived hCiPSCs could form cells comprising the three germ layers *in vivo* (Figure 2H).

To examine the genomic integrity of the hADSC-derived hCiPSCs, we performed high-resolution G-banding analysis, which confirmed that all 12 hCiPSC lines had normal diploid karyotypes (Figures 2I and S1E). Additionally, DNA fingerprinting analyses showed that all hCiPSCs were derived from their parental cells (Table S1). Taken together, these data demonstrated that hADSC-derived hCiPSCs are similar to hESCs in terms of gene expression and differentiation potential, and they can maintain genetic integrity *in vitro*.

### Differentiation of hCiPSCs into the RPE lineage

To induce hCiPSC differentiation into the RPE lineage, we used the differentiation protocol reported by Ye et al^**17**^. Because previous studies revealed that RPE progenitor formation is a critical step in the generation of RPE cells, we analyzed the expression patterns of PAX6 and MITF, key transcription factors modulating RPE progenitor induction^**18**^, on day 12 after initiation of differentiation. Notably, more than 90% of the differentiated cells strongly co-expressed PAX6 and MITF (Figure 3A), suggesting highly efficient induction of RPE progenitor cells from hCiPSCs. To further explore the differentiation of hCiPSCs into the RPE lineage, we performed RT-qPCR analysis that assessed dynamic changes in pluripotency marker genes and RPE marker genes (Figure 3B and S2A). Consistent with the immunofluorescence data, we found that the expression levels of multiple RPE marker genes (e.g., *TYRP1, PEDF, PAX6, MITF, RPE65*, and *BEST1*) gradually increased during differentiation (Figure 3B and S2A). These results suggested that RPE progenitors could be efficiently induced from hADSC-derived hCiPSCs. Furthermore, 5 weeks after differentiation, pigmented colonies with RPE appearance were evident, as previously described^**17**^ (Figure 3C).

**Figure 3.**
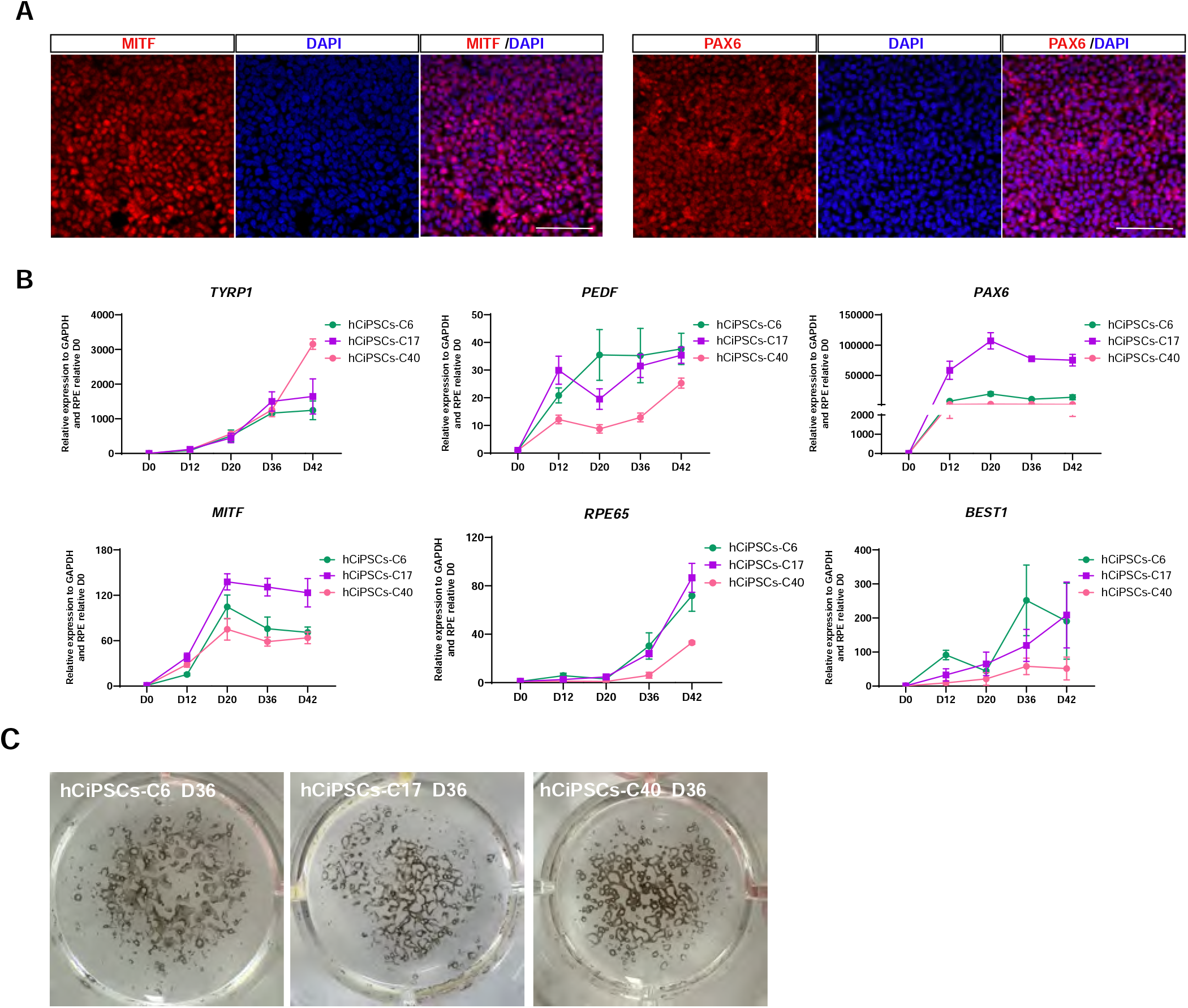
Differentiation of RPE cells from hCiPSCs. (A) Immunostaining for PAX6 and MITF in RPE progenitors on day 12. Scale bar, 100 μm. (B) RT-qPCR analysis of RPE marker genes throughout the differentiation process. Values are presented as means ± SDs; n = 3. (C) Macroscopic photographs of pigmented cells differentiated from various hCiPSC lines on day 36, shown in 12-well plates.

To generate pigmented RPE cells, we cultured RPE progenitor cells in RPE maturation medium (Figure S2B). After 13 days, pigment cells could be observed and on the day 20, RPE pigmentation significantly gathered as evidenced by the emergence of pigmented hexagonal cells (Figure 4A). To examine the characteristics of these cells, we performed immunostaining analysis on day 20 after initiation of induction; the results revealed that distinct RPE markers (e.g., BEST1 and ZO1) were strongly expressed (Figure 4B). RT-qPCR analysis showed the expression of RPE marker genes, such as *BEST1, TRP2, OTX2, PEDF, RPE65*, and *TYRP1* (Figure 4C).

**Figure 4.**
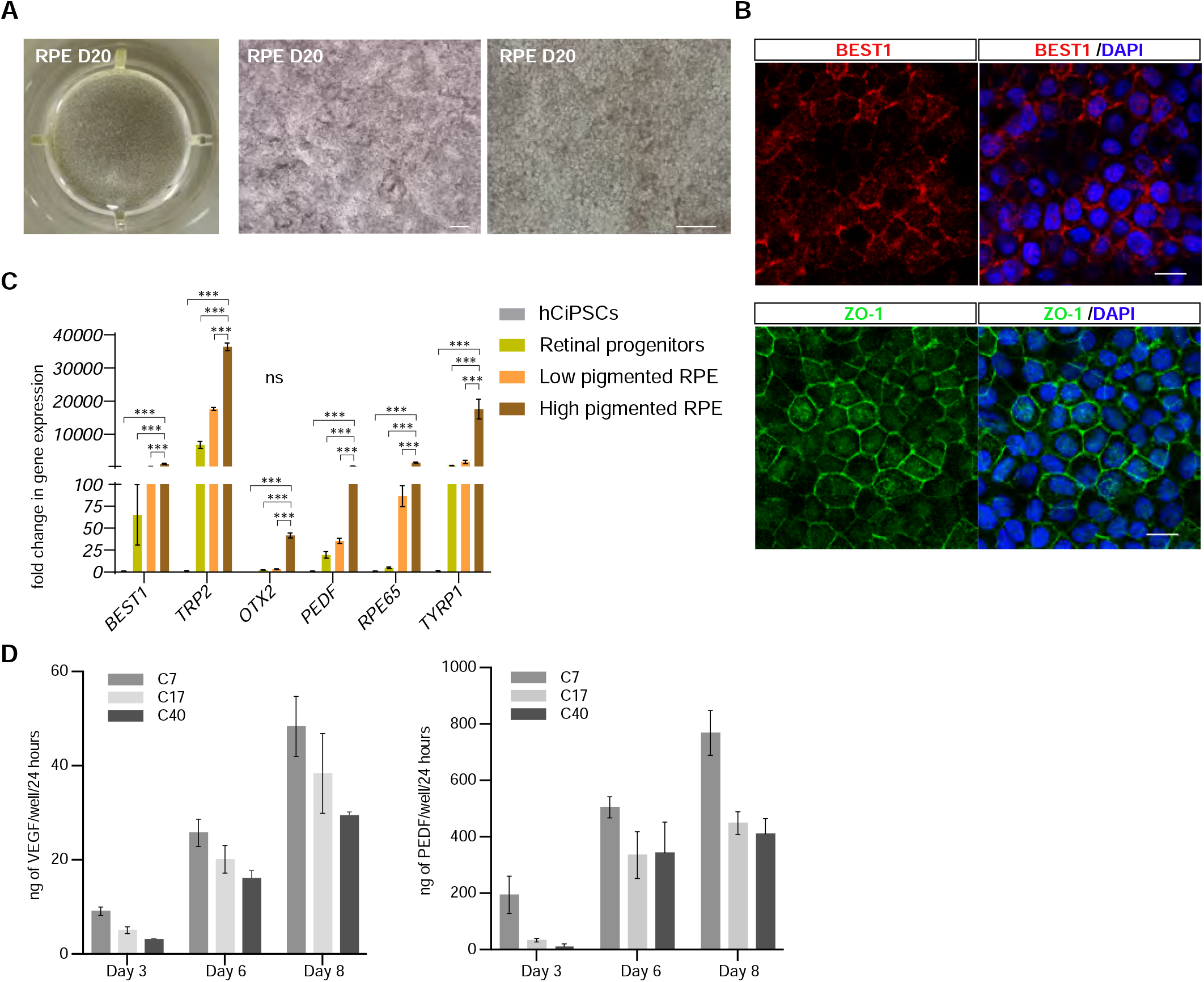
Characterization of mature RPE cells differentiated from hCiPSCs. (A) Macroscopic photographs and phase-contrast images of mature RPE cells. Scale bar, 100μm (left) and 50μm (right) . (B) Immunostaining for BEST1 and ZO1 in RPE cells on day 60. Scale bar, 10 μm. (C) RT-qPCR analysis of the expression of mature RPE markers in hCiPSC-derived mature RPE cells. Values are presented as means ± SDs; n = 3. (D) VEGF (left) and PEDF (right) secretion from hCiPSC-derived mature RPE cells, as measured by ELISAs. Values are presented as means ± SDs; n = 3.

Finally, we analyzed the functions of hCiPSC-derived RPE cells. We utilized enzyme-linked immunosorbent assays (ELISAs) to measure the secretion of pigment epithelium-derived factor (PEDF) and vascular endothelial growth factor (VEGF), which are critical for RPE-mediated maintenance of retinal function. High levels of PEDF and VEGF were detected in the medium of cultured hCiPSC-derived RPE cells; and with the increase of cultivation time, there is a noticeable increase with the levels of PEDF and VEGF. (Figure 4D). Overall, these results demonstrated that functional RPE cells could be efficiently generated from hCiPSCs.

## Discussion

In the present study, we reprogrammed hADSCs into hCiPSCs through a three-step induction process following a detailed protocol^**14**^ and the commercially available Human iPS Cell Chemical Reprogramming Kit (BeiCell™). In total, 12 hCiPSC lines were generated from hADSCs; these cell lines were indistinguishable from hESCs in terms of morphologies, gene expression patterns, and differentiation potentials. Moreover, hCiPSCs could be efficiently induced to differentiate into RPE progenitor cells and then RPE cells through a stepwise differentiation protocol. Our findings demonstrated the efficient induction of RPE cells from hCiPSCs, highlighting the potential for use of clinical-grade autologous hCiPSCs in eye disease treatment.

Compared with the genetic approach, chemical induction represents a simple and robust method for generation of hiPSCs. By simply changing the chemical reprogramming medium, primary hCiPSC colonies emerged within 32 days; this method is particularly valuable for researchers with limited experience in genetic manipulation or cell fate reprogramming. Moreover, consistent with recent reports concerning efficient generation of hCiPSCs from human somatic cells^**12**^, we successfully generated 40 hCiPSC lines. These lines provided sufficient resources for subsequent evaluation and identification of cell lines with RPE differentiation potential. Overall, our results suggest that the chemical reprogramming approach is feasible for generating hiPSCs that can be used in subsequent applications; this approach is valuable for autologous applications involving hiPSCs.

Our results also demonstrated efficient differentiation of RPE cells from hCiPSCs through a two-dimensional differentiation protocol. Compared with three-dimensional differentiation protocols, two-dimensional differentiation protocols are generally simpler and allow for greater control, leading to higher differentiation efficiency^**19**^. In our study, we used a two-dimensional differentiation protocol developed by Ye et al.^**17**^ to generate RPE cells from hCiPSCs; this protocol utilized small molecules to accelerate differentiation, rather than the conventional method of spontaneous differentiation. Notably, the appearance of pigmentation in hCiPSC derivatives was evident as early as day 23 after initiation of differentiation. At around 30 days after initiation of differentiation, significant pigmentation was visible in almost all of the 12 cell lines tested. These results are consistent with the initial reports of RPE cell differentiation from hiPSCs using this protocol and substantially shorter than the duration of spontaneous differentiation (i.e., 3 to 8 months)^**17,20–24**^. Additionally, the secretion of PEDF and VEGF suggests that hCiPSC-derived RPE cells exhibit functional maturity. These results indicate a possible new route for the generation of sufficient autologous functional RPE cells that can be used in clinical treatment of retinal diseases.

To achieve the full potential of hCiPSC-based autologous therapies for retinal diseases, two major issues must be addressed. First, a thorough evaluation of the genomic stability and safety of autologous hiPSCs and their retinal derivatives should be performed prior to clinical use, because recent studies have revealed that acquired cancer-related mutations are prevalent in numerous hiPSC lines and their derivatives^**25,26**^. Indeed, the genetic instability evident in hiPSCs and their RPE derivatives was a key factor leading to the suspension of the AMD clinical trial at RIKEN^**10**^. It is possible that the genetic factors utilized in hiPSC generation may have contributed to this phenomenon, as evidenced by their upregulated expression in various tumor types^**27–29**^. In contrast, chemical induction avoids the use of genetic factors; therefore, future studies should explore whether hCiPSCs could bypass the potential safety issues associated with overexpression of genetic factors. Second, considering the complex, multilayered structure of the retina and the involvement of its diverse cell layers in disease progression, exclusive reliance on the replacement of dysfunctional RPE cells may not achieve optimal therapeutic effects^**30,31**^. Therefore, future studies should explore alternative hCiPSC-based approaches such as co-transplantation of RPE cells with retinal progenitor cells derived from hCiPSCs, as well as the transplantation of more complex and structured hCiPSC-derived multilayered retinal cells or organoids^**32**^.

In summary, our study effectively utilized a chemical reprogramming method to generate hCiPSCs from human somatic cells, then differentiated the hCiPSCs into RPE cells. Chemical induction has considerable promise in overcoming the limitations associated with iPSC reprogramming via genetic techniques. This non-genetic method has the potential to generate autologous iPSCs for future clinical applications in a reproducible, controlled, user-friendly, and cost-effective manner.

## Methods and Materials

### Isolation and culture of human adipose tissue-derived stromal cells in vitro

hADSCs were obtained from donated fat tissue, and the informed consent has been obtained. Under sterile conditions, adipose tissue (2∼4 cm^3^) was washed twice with PBS solution (AQ) containing 2% penicillin-streptomycin (Invitrogen), and then cut to 1∼2 mm^3^ with scissors. To dissociate the fat tissue, add 5∼10ml collagenase IV (ThermoFisher) solution (2mg/ml) in 100mm Petri dish and then incubate at 37 °C for 1 hour. After incubation, 10∼20 ml of high-glucose DMEM (gibco) supplemented with 10% fetal bovine serum (FBS) (gibco) and 1% penicillin-streptomycin (ThermoFisher) were added. After ADSCs were fully dissociated, the suspension was collected into two 50ml centrifuge tubes, diluted with 10% FBS-DMEM medium to 30∼40ml in each tube, and then shaken for 1∼2 min to fully release the cells. The suspension was centrifuged at 400*g* for 5 min. After removing the supernatant, the cells were resuspended in mesenchymal stem cell growth medium (MSCGM) (Promo cell), and then inoculated with a suitable Petri dish according to the amounts of cells. The next day, fresh MSCGM was replaced to remove non-adherent cells. hADSCs were fully integrated within 3∼5 days and ready for reprogramming to hCiPSCs.

### Reprogramming of hADSCs to hCiPSCs

To generate hCiPSCs from hADSCs, we followed a detailed protocol^**14**^ and utilized Human iPS Cell Chemical Reprogramming Kit (BeiCell™) to reprogram the hADSCs (passage 2∼4) through a three-step induction. It was suitable to inoculate 12 well plates with 1×10^4^ cells per well. The hADSCs were cultured in DMEM-F12 medium (gibco) with 15% FBS (gibco) for 1 day, and then switched to chemical induction medium. The first stage of induction lasted 8 days. Medium A was used until the confluence of the monolayered epithelial cells was close to 100%. Medium B was used for 12 days at the second stage of induction, and by the end of this stage, a large number of multi-layer clones emerged. The last stage used medium C, D, E in turn and lasted 12 days with the medium changed every 4 days. At the end of the induction stage, we selected the wells in which the clones were suitable for passage. The cells in the selected wells were digested with prewarmed enzyme Accutase (Merck), and cultured with medium F on the laminin 521 (stemcell)-coated plate. The passage ratio was 1:12, and medium F was changed every three days. Generally, the clone of hCiPSCs enlarged after six days, then the culture medium was changed to mTeSR™ Plus (stemcell). When the homogeneous clones with typical morphology appear, clear edges could be observed under the microscope. Cell clones were collected to further build cell lines. 0.3 ml ReLeSR was added to each well of the 12 well plate and digested at room temperature for 2 min, and 37 °C for 2∼3 min. The clone was scraped off carefully with a glass needle, and was absorbed with 20μl medium. The selected clones were inoculated on the Matrigel (Corning) -coated culture plate and cultured using mTeSR™ Plus medium.

### RPE differentiation

In accordance with Ye’s protocol^**17**^, the directed differentiation of hCiPSCs into RPE cells was achieved within 36 days. hCiPSCs were treated with Accutase (Merck) for 5 minutes to dissociate into single cells, and then plated onto Martrigel (Corning) -coated 12-well culture plates. It is advisable to have 4.0×10^4^ cells per well. For days 0∼17 of differentiation, hCiPSCs were treated with IMDM/Ham’s F12 medium (1:1) (both were from gibco) containing 10 μM Y-27632 (Selleck), 10% KnockOut Serum Replacement (gibco), 1% Chemically Defined Lipid Concentrate (gibco), and 2 mM L-Glutamine (gibco). 100 nM LDN193189 (MCE), 500 nM A-83-01 (Selleck), 1 μM IWR-1-endo (MCE) were added for the initial 6 days. 3 μM CHIR99021 (MCE) and 2 μM SU5402 (MCE) were added for days 6∼11. 10 mM nicotinamide (Merck) were added additionally for days 12∼17 in IMDM/F12. From day 18∼23, the medium was changed to DMEM/F12 (gibco) containing 10% KnockOut Serum Replacement (gibco), 1% N2 supplement (gibco), 2 mM Glutamax (gibco) and 10 mM nicotinamide (Merck). After 24 days, for the further maturation of hCiPSCs-RPE, RPE maintenance medium containing 67% high-glucose DMEM (gibco), 29% Ham’s F12 (gibco), 2% B27 supplement minus vitamin A (gibco), 2 mM Glutamax (gibco) were used. Culture medium was changed every day. After the appearance of obvious pigment clumps (between 35∼40 days), we mechanically separated the pigment cell areas using glass needles. Subsequently, the pigment cells were digested into single cells using 0.25% Trypsin-EDTA for 15 minutes, and then replated on culture plate coated with Matrigel (Corning). RPE growing medium (Ham’s F12 (gibco), 10% FBS (gibco), 2 mM L-Glutamine (gibco)), and 1% penicillin-streptomycin (Invitrogen)) was used for cell culture. After 14 days, the medium was changed to RPE maintenance medium with 10 ng/mL basic fibroblast growth factor (R&D Systems) and 0.5 μM SB431542 (MCE).

### Embryoid body formation

For EBs formation, hCiPS cells were digested as small clumps by ReLeSR and cultured in ultra-low attachment 6-well cell culture plates in mTeSR™ Plus Medium for 1 day, and then transformed into differentiation medium for 16 days. The differentiation medium was consisted of high-glucose DMEM supplemented with 20% FBS and 1% penicillin/streptomycin. EBs were then collected and attached into 6-well cell culture plates with pre-incubated Matrigel for 6 days in the same medium, fixed and detected using immunostaining analysis. Medium was changed every day.

### Teratoma formation

For teratoma formation, approximately 2 × 10^6^ hCiPSCs cells were collected by ReLeSR, resuspended in pre-cooled 200 µl Matrigel and subcutaneously injected to the hind limbs of male immunodeficient NPG mice (Vitalstar Biotechnology, Beijing) aged 2-3 months. Tumor formation occurs 6-8 weeks later. When the size of the teratoma approached 1∼1.5 cm, it can be retrieved and then embedded in paraffin. All animal experiments were approved by the Institutional Animal Care and Use Committee of Peking University and performed according to the Animal Protection Guidelines of Peking University.

### RT–qPCR analysis

Total RNA was isolated using the Direct-zol RNA MiniPrep Kit from Zymo Research. cDNA was synthesized 1.5 μg of total RNA using TransScript First-Strand cDNA Synthesis SuperMix (TransGen Biotech). Real-time PCR was performed with the 2× S6 SYBR Premix EsTaq plus (Scintol) on the CFX Connect Real-Time System (Bio-Rad). The data were analysed using the 2^-ΔΔCt^ method. *GAPDH* was used as a control to normalize the expression of target genes. The primer pairs used in this study are described in Table S2.

### Immunofluorescence

For immunofluorescence staining, cells were fixed for 30 min in 4% paraformaldehyde (DingGuo) at room temperature, and then blocked by PBS containing 0.1% Triton™ X-100 (Sigma-Aldrich) and 2% donkey serum (Jackson) at 37 °C for 1 hour. Primary antibodies incubation with appropriate dilutions were incubated overnight at 4°C in the same buffer. After washing three times with DPBS, the secondary antibodies were incubated in PBS containing 2% donkey serum for 1 hour at room temperature. DNA was stained with DAPI solution (Roche). Antibody details are provided in Table S3.

### Karyotype analysis

The karyotype (chromosomal G-band) analysis were contracted out at Beijing Jiaen Hospital, using standard protocols for high-resolution G-banding (400G–500G) and analysed by CytoVision (Leica). For each analysis, the number of chromosomes as well as the presence of structural chromosomal abnormalities of at least 20 metaphases were examined.

### STR analysis

Short-tandem repeat (STR) analysis was contracted out at Beijing Microread Genetics. In brief, the genomic DNA was amplified by PCR using the STR Multi-amplification Kit (Microreader 21 ID System) and analysed using the ABI 3730xl DNA Analyzer (Applied Biosystems), the data were analyzed using GeneMapperID-X software. Twenty-one loci were analyzed for each sample and no cross-contamination of other cell line was found.

### Population doubling time

The growth rate was determined by a haemocytometer count to calculate the number of cells as a function of time. The doubling time (DT) was calculated following the formula: DT=t/[log2/(logN_t_−logN_0_)], where N_t_ is the number of cells at time t and N_0_ is the number of cells at time zero.

### RNA sequencing (RNA-seq) and data analysis

Total RNA was isolated by using Direct-zol RNA Mini PrepKit (ZymoResearch), and checked for integrity using an Agilent Bioanalyzer 2100. mRNA is enriched by oligo (dT) -attached magnetic beads from total RNA, strand specific transcriptome library is constructed, sequenced by DNBSEQ high-throughput platform, all the above processes were performed at BGI Technology.

The sequencing data was filtered with SOAPnuke^**33**^, afterwards clean reads were obtained and stored in FASTQ format. The subsequent analysis and data mining were performed on Dr. Tom Multi-omics Data mining system (https://biosys.bgi.com).

Expression level of gene was calculated by RSEM (v1.3.1)^**34**^. The heatmap was drawn by pheatmap (v1.0.8)^**35**^ according to the gene expression difference. Essentially, differential expression analysis was performed using the DESeq2(v1.4.5)^**36**^ with Q value ≤ 0.05.

### Enzyme-linked immunosorbent assay

The culture media were collected following a 24-hour exposure to the cells after 30 days of enrichment. The levels of human PEDF and VEGF were determined using Human VEGF ELISA Kit (Abcam) and Human PEDF ELISA Kit (Abcam). The plate growth area and media volume were taken into account when calculating the levels of PEDF and VEGF, expressed in ng/mm^2^.

### Statistical analysis

All values are shown as means ± SD. The number of biological replicates and the method for statistical analysis and n values are reported in the figures. Statistical analysis was performed in GraphPad Prism (version 8).

## References

1. Soldner F, Jaenisch R. Stem Cells, Genome Editing, and the Path to Translational Medicine. Cell. 2018;175(3):615–632. doi:10.1016/j.cell.2018.09.010

2. Rowe RG, Daley GQ. Induced pluripotent stem cells in disease modelling and drug discovery. Nat Rev Genet. 2019;20(7):377–388. doi:10.1038/s41576-019-0100-z

3. Yamanaka S. Pluripotent Stem Cell-Based Cell Therapy-Promise and Challenges. Cell Stem Cell. 2020;27(4):523–531. doi:10.1016/j.stem.2020.09.014

4. Kashani AH, Lebkowski JS, Rahhal FM, et al. A bioengineered retinal pigment epithelial monolayer for advanced, dry age-related macular degeneration. Sci Transl Med. 2018;10(435):eaao4097. doi:10.1126/scitranslmed.aao4097

5. Schwartz SD, Hubschman JP, Heilwell G, et al. Embryonic stem cell trials for macular degeneration: a preliminary report. Lancet. 2012;379(9817):713–720. doi:10.1016/S0140-6736(12)60028-2

6. Schwartz SD, Regillo CD, Lam BL, et al. Human embryonic stem cell-derived retinal pigment epithelium in patients with age-related macular degeneration and Stargardt’s macular dystrophy: follow-up of two open-label phase 1/2 studies. Lancet. 2015;385(9967):509–516. doi:10.1016/S0140-6736(14)61376-3

7. Mandai M, Kurimoto Y, Takahashi M. Autologous Induced Stem-Cell-Derived Retinal Cells for Macular Degeneration. N Engl J Med. 2017;377(8):792–793. doi:10.1056/NEJMc1706274

8. da Cruz L, Fynes K, Georgiadis O, et al. Phase 1 clinical study of an embryonic stem cell-derived retinal pigment epithelium patch in age-related macular degeneration. Nat Biotechnol. 2018;36(4):328–337. doi:10.1038/nbt.4114

9. Kobold S, Bultjer N, Stacey G, Mueller SC, Kurtz A, Mah N. History and current status of clinical studies using human pluripotent stem cells. Stem Cell Reports. 2023;18(8):1592–1598. doi:10.1016/j.stemcr.2023.03.005

10. Garber K. RIKEN suspends first clinical trial involving induced pluripotent stem cells. Nat Biotechnol. 2015;33(9):890–891. doi:10.1038/nbt0915-890

11. Neofytou E, O’Brien CG, Couture LA, Wu JC. Hurdles to clinical translation of human induced pluripotent stem cells. J Clin Invest. 2015;125(7):2551–2557. doi:10.1172/JCI80575

12. Liuyang S, Wang G, Wang Y, et al. Highly efficient and rapid generation of human pluripotent stem cells by chemical reprogramming. Cell Stem Cell. 2023;30(4):450–459.e9. doi:10.1016/j.stem.2023.02.008

13. Hou P, Li Y, Zhang X, et al. Pluripotent stem cells induced from mouse somatic cells by small-molecule compounds. Science. 2013;341(6146):651–654. doi:10.1126/science.1239278

14. Guan J, Wang G, Wang J, et al. Chemical reprogramming of human somatic cells to pluripotent stem cells. Nature. 2022;605(7909):325–331. doi:10.1038/s41586-022-04593-5

15. Zhang M, Wang L, An K, et al. Lower genomic stability of induced pluripotent stem cells reflects increased non-homologous end joining. Cancer Commun (Lond). 2018;38(1):49. doi:10.1186/s40880-018-0313-0

16. Shyh-Chang N, Zhu H, Yvanka de Soysa T, et al. Lin28 enhances tissue repair by reprogramming cellular metabolism. Cell. 2013;155(4):778–792. doi:10.1016/j.cell.2013.09.059

17. Ye K, Takemoto Y, Ito A, et al. Reproducible production and image-based quality evaluation of retinal pigment epithelium sheets from human induced pluripotent stem cells. Sci Rep. 2020;10(1):14387. doi:10.1038/s41598-020-70979-y

18. Diacou R, Nandigrami P, Fiser A, Liu W, Ashery-Padan R, Cvekl A. Cell fate decisions, transcription factors and signaling during early retinal development. Progress in Retinal and Eye Research. 2022;91:101093. doi:10.1016/j.preteyeres.2022.101093

19. Plaza Reyes A, Petrus-Reurer S, Padrell Sánchez S, et al. Identification of cell surface markers and establishment of monolayer differentiation to retinal pigment epithelial cells. Nat Commun. 2020;11(1):1609. doi:10.1038/s41467-020-15326-5

20. Choudhary P, Booth H, Gutteridge A, et al. Directing Differentiation of Pluripotent Stem Cells Toward Retinal Pigment Epithelium Lineage. Stem Cells Transl Med. 2017;6(2):490–501. doi:10.5966/sctm.2016-0088

21. Kuroda T, Ando S, Takeno Y, Kishino A, Kimura T. Robust induction of retinal pigment epithelium cells from human induced pluripotent stem cells by inhibiting FGF/MAPK signaling. Stem Cell Res. 2019;39:101514. doi:10.1016/j.scr.2019.101514

22. Lange L, Esteban MA, Schambach A. Back to pluripotency: fully chemically induced reboot of human somatic cells. Signal Transduct Target Ther. 2022;7(1):244. doi:10.1038/s41392-022-01109-5

23. Buchholz DE, Hikita ST, Rowland TJ, et al. Derivation of functional retinal pigmented epithelium from induced pluripotent stem cells. Stem Cells. 2009;27(10):2427–2434. doi:10.1002/stem.189

24. Maruotti J, Sripathi SR, Bharti K, et al. Small-molecule-directed, efficient generation of retinal pigment epithelium from human pluripotent stem cells. Proc Natl Acad Sci U S A. 2015;112(35):10950–10955. doi:10.1073/pnas.1422818112

25. Rouhani FJ, Zou X, Danecek P, et al. Substantial somatic genomic variation and selection for BCOR mutations in human induced pluripotent stem cells. Nat Genet. 2022;54(9):1406–1416. doi:10.1038/s41588-022-01147-3

26. Lezmi E, Jung J, Benvenisty N. High prevalence of acquired cancer-related mutations in 146 human pluripotent stem cell lines and their differentiated derivatives. Nat Biotechnol. Published online January 9, 2024. doi:10.1038/s41587-023-02090-2

27. Dang CV. MYC on the path to cancer. Cell. 2012;149(1):22–35. doi:10.1016/j.cell.2012.03.003

28. Wang YJ, Herlyn M. The emerging roles of Oct4 in tumor-initiating cells. Am J Physiol Cell Physiol. 2015;309(11):C709–718. doi:10.1152/ajpcell.00212.2015

29. Ding LN, Yu YY, Ma CJ, Lei CJ, Zhang HB. SOX2-associated signaling pathways regulate biological phenotypes of cancers. Biomed Pharmacother. 2023;160:114336. doi:10.1016/j.biopha.2023.114336

30. McLelland BT, Lin B, Mathur A, et al. Transplanted hESC-Derived Retina Organoid Sheets Differentiate, Integrate, and Improve Visual Function in Retinal Degenerate Rats. Invest Ophthalmol Vis Sci. 2018;59(6):2586–2603. doi:10.1167/iovs.17-23646

31. Barnea-Cramer AO, Wang W, Lu SJ, et al. Function of human pluripotent stem cell-derived photoreceptor progenitors in blind mice. Sci Rep. 2016;6:29784. doi:10.1038/srep29784

32. Hirami Y, Mandai M, Sugita S, et al. Safety and stable survival of stem-cell-derived retinal organoid for 2 years in patients with retinitis pigmentosa. Cell Stem Cell. 2023;30(12):1585-1596.e6. doi:10.1016/j.stem.2023.11.004

33. Li R, Li Y, Kristiansen K, Wang J. SOAP: short oligonucleotide alignment program. Bioinformatics. 2008;24(5):713–714. doi:10.1093/bioinformatics/btn025

34. Li, B. & Dewey, C. N. RSEM: accurate transcript quantification from RNA-Seq data with or without a reference genome.BMC Bioinformatics 2011;12:323. doi:10.1186/1471-2105-12-323.

35. Raivo Kolde. Package ‘pheatmap’. 2019-01-04 13:50:12 UTC.

36. Love, M. I., Huber, W. & Anders, S. Moderated estimation of fold change and dispersion for RNA-seq data with DESeq2. Genome Biol. 2014; 15(121):550. doi:10.1186/s13059-014-0550-8.

