## Supplementary Figure for "Efficient differentiation of human retinal pigment epithelium cells from chemically induced pluripotent stem cells"

### Figure S1

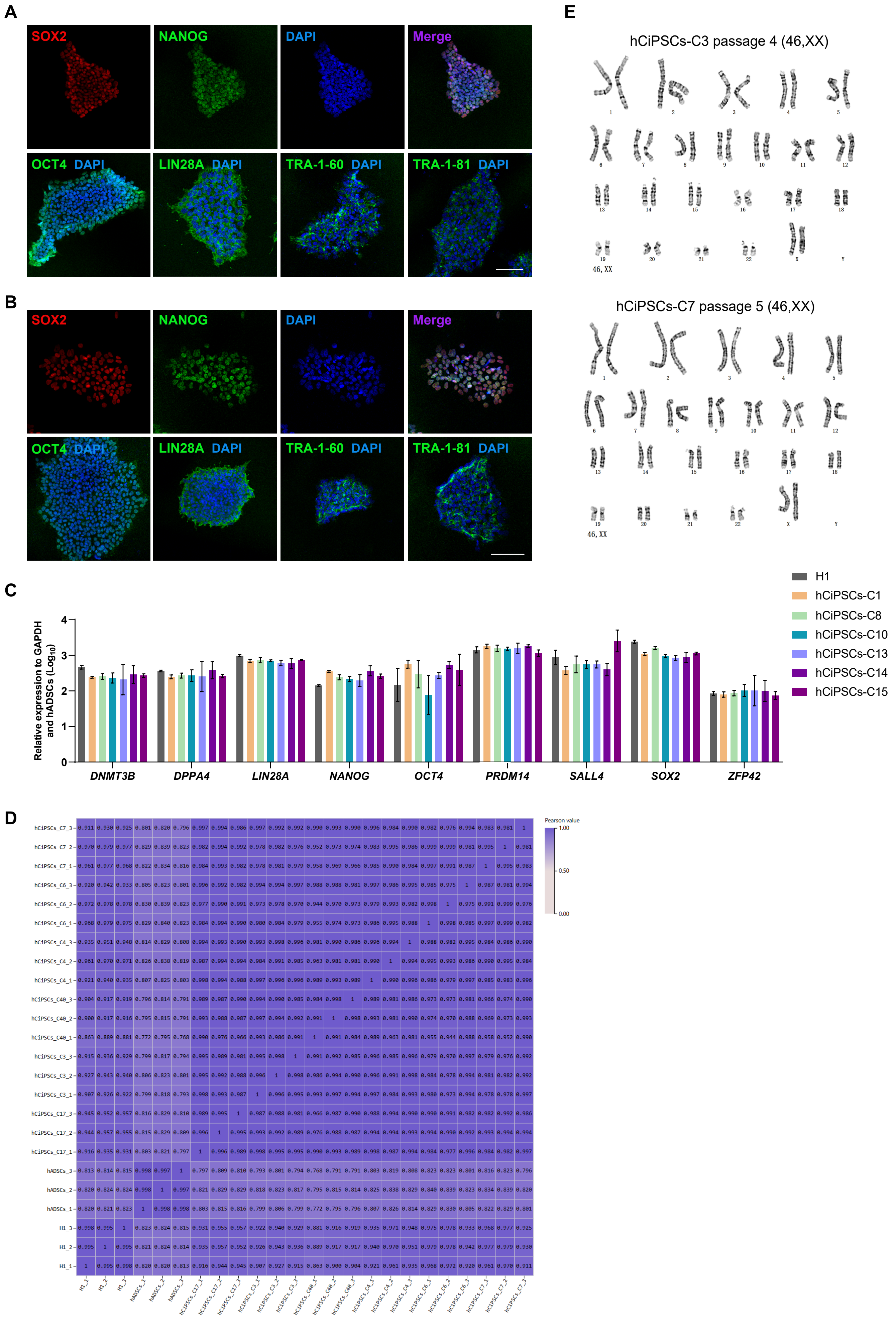
A
SOX2
NANOG
DAPI
Merge
OCT4 DAPI
LIN28A DAPI
TRA-1-60 DAPI
TRA-1-81 DAPI
B
SOX2
NANOG
DAPI
Merge
OCT4 DAPI
LIN28A DAPI
TRA-1-60 DAPI
TRA-1-81 DAPI
C
Relative expression to GAPDH and hADSCs (Log10)
DNMT3B
DPPA4
LIN28A
NANOG
OCT4
PRDM14
SALL4
SOX2
ZFP42
H1
hCiPSCs-C1
hCiPSCs-C8
hCiPSCs-C10
hCiPSCs-C13
hCiPSCs-C14
hCiPSCs-C15
D
Pearson value
E
hCiPSCs-C3 passage 4 (46,XX)
46, XX
hCiPSCs-C7 passage 5 (46,XX)
46, XX

**Figure S1. Characterization of hADSC-derived hCiPSCs.** (A-B) Immunofluorescence staining for pluripotency markers in hCiPSCs-C3 (A) and hCiPSCs-C7 (B). Scale bar, 100 μm. (C) Relative expression levels of pluripotency marker genes in hESCs (H1) and hADSC-derived hCiPSCs, as determined by RT-qPCR. Data are presented as means ± SDs; n = 3. (D) Correlation analysis of the global transcriptomes of H1, hADSCs, and hADSC-derived hCiPSCs. (E) Karyotype analysis showing hADSC-derived hCiPSCs-C3 and hCiPSCs-C7 with normal diploid chromosomal content.

#### Figure S2

A

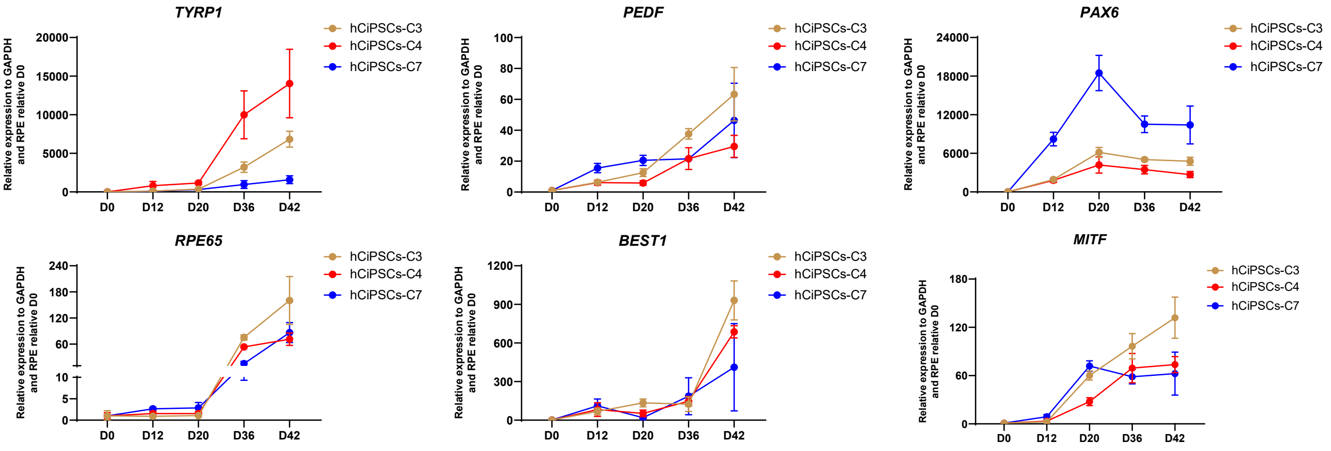
TYRP1
PEDF
PAX6
RPE65
BEST1
MITF
hCiPSCs-C3
hCiPSCs-C4
hCiPSCs-C7
Relative expression to GAPDH and RPE relative D0
D0
D12
D20
D36
D42

B

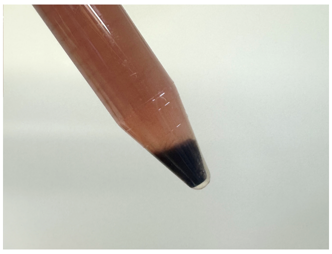

**Figure S2. Characterization of RPE cells derived from other hCiPSCs.** (A) RT-qPCR analysis of RPE marker genes throughout the differentiation process of hCiPSCs-C3/C4/7. Values are presented as means ± SDs; n = 3. (B) Cell pellets after collection and dissociation of hCiPSC-derived RPE cells.
